# NeuroMechanics: Electrophysiological and Computational Methods to Accurately Estimate the Neural Drive to Muscles in Humans *In Vivo*

**DOI:** 10.1101/2024.01.03.574073

**Authors:** Arnault H. Caillet, Andrew T.M. Phillips, Luca Modenese, Dario Farina

**Author notes:** (DF) (LM). These authors have equal contributions and share the senior authorship (Luca Modenese, Dario Farina). **Conflict of interests:**The authors declare no competing financial interests.

## Abstract

The ultimate neural signal for muscle control is the neural drive sent from the spinal cord to muscles. This neural signal comprises the ensemble of action potentials discharged by the active spinal motoneurons, which is transmitted to the innervated muscle fibres to generate forces. Accurately estimating the neural drive to muscles in humans *in vivo* is challenging since it requires the identification of the activity of a sample of motor units (MUs) that is representative of the active MU population. Current electrophysiological recordings usually fail in this task by identifying small MU samples with over-representation of higher-threshold with respect to lower-threshold MUs. Here, we describe recent advances in electrophysiological methods that allow the identification of more representative samples of greater numbers of MUs than previously possible. This is obtained with large and very dense arrays of electromyographic electrodes. Moreover, recently developed computational methods of data augmentation further extend experimental MU samples to infer the activity of the full MU pool. In conclusion, the combination of new electrode technologies and computational modelling allows for an accurate estimate of the neural drive to muscles and opens new perspectives in the study of the neural control of movement and in neural interfacing.

## Introduction

Skeletal muscles are multiscale structures composed of individual units called motor units (MUs). A MU comprises a motoneuron (MN) and the muscle fibres it innervates (Heckman & Enoka, 2012). The neural drive to muscle refers to the ensemble of action potentials discharged by the active MN population and transmitted to the muscle fibres (Farina et al., 2014a). At the muscle-scale, the neural drive is seen as the compound neural control signal from the MN pool, which is received by the macroscopic muscle actuator. Therefore, at the muscle-scale, the neural drive represents the sum of the MNs’ discharge activities, which is commonly mathematically described as the temporal cumulative series of MNs’ discharges. Because the dynamic response of muscles has small bandwidth, the neural drive can control force at frequencies up to 4-to-10 Hz (Farina & Negro, 2015; Enoka & Farina, 2021).

The intensity of the neural drive is determined by the number of active MNs (recruitment) and their firing rates (rate coding). The recruitment and rate coding dynamics of the MN pool are determined by the amplitude of the net excitatory synaptic drive the MNs receive and on the intrinsic excitability of the MNs, which is influenced by neuromodulatory inputs (Heckman & Enoka, 2012; Binder et al., 2020; Avrillon et al., 2023a). In response to a rising net excitatory synaptic drive, the MUs are recruited sequentially in order of size (Henneman’s size principle; Henneman, 1957; Henneman, 1981), starting with the smallest MUs (small MNs, low current and force thresholds for recruitment, low generated forces). Human MU pools include many low-threshold MUs and few high-threshold MUs following an exponential continuous distribution (Duchateau & Enoka, 2022; Caillet et al., 2023a). The smaller MUs usually attain higher discharge frequencies (De Luca & Hostage, 2010) and undergo a more rapid acceleration of the firing rate at recruitment and a more pronounced rate limiting than larger MUs (Avrillon et al., 2023a).

The neural drive to a muscle determines the force the muscle produces. The muscle fibres build cellular force based on the frequency of discharged action potentials they receive from their innervating active MNs (Heckman & Enoka, 2004). The muscle aggregates the force-generating dynamics of the active MUs into a macroscopic pulling muscle force that is transmitted to the skeletal system. By modulating the recruitment and rate coding dynamics of the MN pool, the central nervous system voluntarily controls the muscle force (Enoka & Duchateau, 2017). For these reasons, accurately estimating the neural drive to muscle during human muscle contractions allows for the neuromechanical investigations of neural control strategy of movement, where neural control signals are associated to muscle forces and behaviour (e.g., Hug et al., 2023, Powers & Heckman, 2017; Binder et al., 2020), as well as the establishment of human-machine interfacing (Farina et al., 2017; Farina et al., 2021; Jung et al., 2021). Accurately estimating the neural drive to muscle in humans *in vivo* is however challenged by the difficulty of identifying the firing activity of full MU pools in humans *in vivo* (Figure 1, (Farina & Holobar, 2016; Avrillon et al., 2023a)).

**Figure 1:**
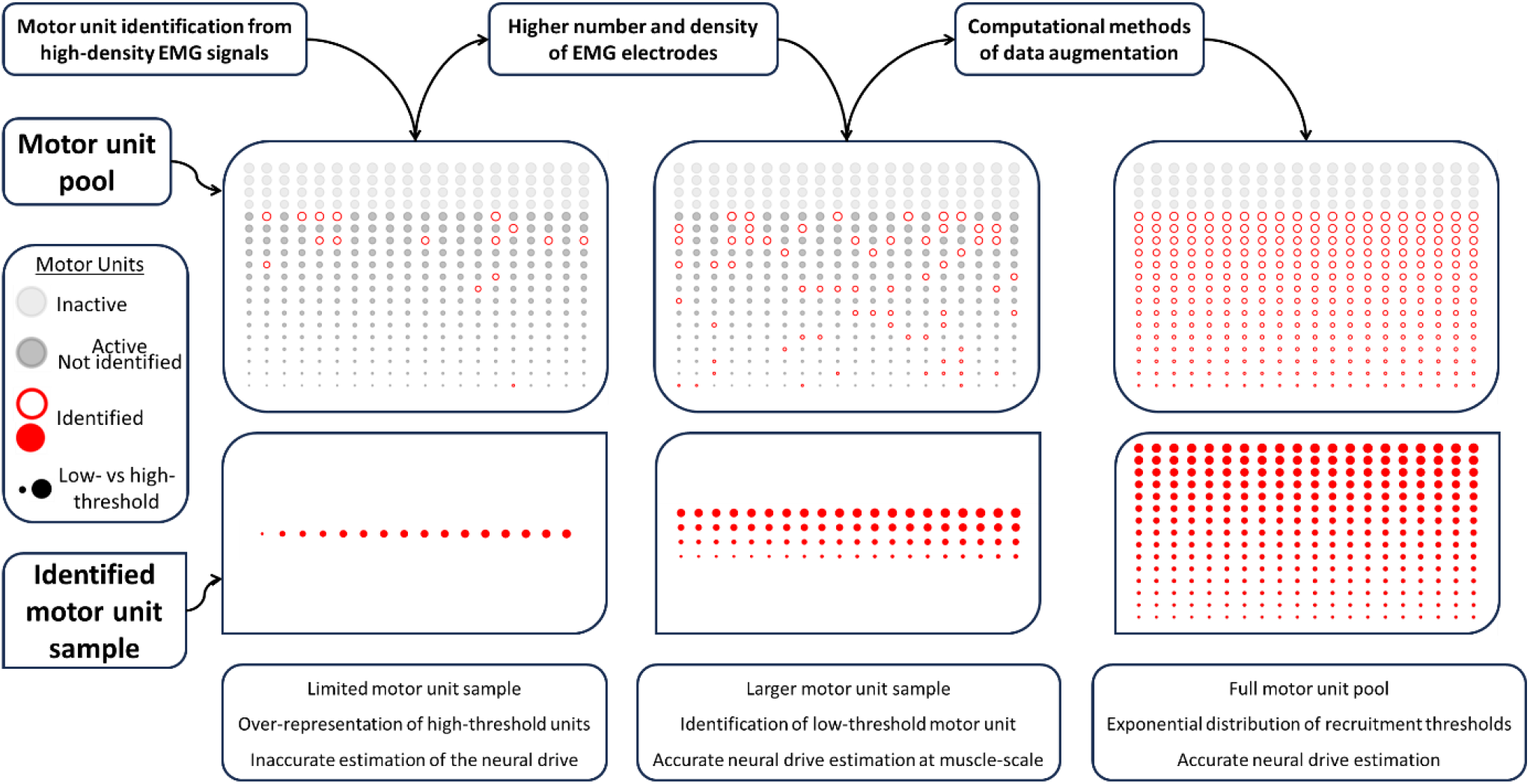
Identification of the discharge activity of the active motor unit (MU) pool with electrophysiological and computational methods to estimate the neural drive to muscle in humans in *vivo*. The subplots in the first row display an exemplary pool of 400 MUs. Each MU is represented by a circle with a size schematically representing the MN size and therefore the MU’s force recruitment threshold, following a physiological exponential distribution across the MU pool (given in Figure 2). The MU population is here ranked from left to right and bottom to top in increasing order of recruitment thresholds. Here, 75% of the MU pool is recruited and 25% of the MUs are not active (first row: light grey circles). A sample of active MUs and their discharge activity (first row: red hollow circles; second row: plain red circles) can be identified in humans *in vivo* from the decomposition of high-density EMG signals. The MU samples commonly identified in this way (first column) are usually small and over-represent high-threshold MUs (large diameter circles). Therefore, these MU samples are usually not representative of the MU pool. Increasing the number and the density of EMG electrodes (second column) allows the identification of more MUs, especially low-threshold MUs, yielding MU samples that are more representative and that provide a more accurate estimation of the neural drive to muscle at the muscle-scale. Computational methods of data augmentation allow the estimation of the discharge behaviour of the MUs not identified experimentally (dark grey plain circles) and fully describe the neural drive to muscle. Adapted from data in the tibialis anterior muscle at 30% maximum voluntary contraction force by Caillet et al. (2023a, 2023b).

This review presents modern electrophysiological and computational methods that allow us to accurately estimate the neural drive to muscle, i.e., the discharge activity of the full MU pool, at least in some muscles and in constrained laboratory conditions. First, we describe electrophysiological methods of surface and intramuscular high-density electromyography (EMG) acquisition and decomposition for identifying experimental samples of MUs that are representative of the full MU pool. The latter condition implies that a sufficient number of MUs is identified and that the identified MUs have recruitment thresholds that follow a physiological exponential distribution, with more low- threshold units than high-threshold units (Figure 1, second column). Then, we review computational methods of data augmentation that estimate the discharge activity of the full MU pool from limited experimental samples of discharging MUs (Figure 1, third column). Finally, we apply these electrophysiological and computational methods to representatively demonstrate the accurate control of a MN-driven neuromuscular model of a MU pool.

## Properties of MNs and MUs and representative MU samples

It is currently not possible to experimentally detect the activity of full pools of active MUs in humans. Therefore, the neural drive to muscle is commonly inferred from limited samples of experimentally observed MUs. These samples must be representative of the active MU pool for accurate estimations of the neural drive. In this section, we will review the distribution of MN and MU properties across the pool, with special emphasis to their interrelations and their contribution to the neural drive. This discussion will be relevant to better specify the issue of representativeness of MU samples and to later explain how data augmentation methods can infer the activity of non-observed MUs from representative samples of observed ones.

### The electrophysiological properties of a MN are inter-related

The input-output function of MNs, i.e., the transformation of net excitatory synaptic current into firing activity, is MN-specific. Indeed, the firing behaviour of a MN depends on its set of intrinsic electrophysiological properties. The MN-specific electrophysiological properties are inter-related with each other for each MN (Henneman, 1981; Burke, 1981; Binder et al., 1996; Powers & Binder, 2001; Kernell, 2006; Heckman & Enoka, 2012), as it can be quantified with mathematical equations that express each property as a function of the others (Table 4 in Caillet et al. (2022a)). According to these relations, the higher the value of current recruitment threshold in a MN, the greater the MN’s capacitance and axonal conduction velocity, while the MN have smaller input resistance, afterhyperpolarization period, and membrane time constant. Besides, recent findings (Avrillon et al., 2023a) show that low-threshold MNs have a higher initial acceleration of firing rates, stronger rate limiting, and a more pronounced hysteresis than high-threshold MNs, because of MN-specific neuromodulatory inputs. Also, low-threshold MNs usually reach higher discharge rates than high-threshold MNs (onion-skin scheme (De Luca & Hostage, 2010)). Combined, these results indicate that the input-output function of MNs is MN-specific and each active MN contributes in a unique way to the neural drive to muscle.

### MN and muscle fibre properties are correlated

The current recruitment threshold and force recruitment threshold of a MU are positively correlated, as indirectly demonstrated from separate measurements of current thresholds, torque recruitment thresholds, twitch torques, and tetanic forces in human and other mammal muscles (Heckman & Enoka, 2012; Caillet et al., 2022a). Furthermore, Caillet et al. (2022a) reported significant correlations between the tetanic force, twitch force, or innervation ratio of a MU and the axonal conduction velocity, input resistance, or current threshold of its innervating MN in cats and rodents. Therefore, the electrophysiological properties of a MN are also correlated with the properties of the muscle fibres it innervates. Notably, the early-recruited low-threshold (in current or force) MUs are activated at low forces before the higher-threshold (in current or force) MUs, that have large number of muscle fibres and therefore produce high forces. MN and MU properties are also quantitively correlated to MN and MU size (Caillet et al., 2022a), so that larger MNs and MUs have higher (current and force) recruitment thresholds (Henneman’s size principle (Henneman, 1981)).

### Distribution of MU thresholds

The inter-related properties of MNs and MUs are distributed continuously across the MU pool’s recruitment range in mammals and do not cluster into discrete groups, as reviewed previously (Heckman & Enoka, 2012) and demonstrated with density histograms in Caillet et al. (2022a) and Duchateau & Enoka (2022) for several MN and MU properties. Of note, these histograms were obtained by merging limited samples of identified MUs from many subjects. This limitation was recently addressed (Avrillon et al., 2023a) when extensive samples of up to ∼200 MUs were identified in single muscles of individual human subjects across the full recruitment range in separate contractions. Figure 2 reports the continuous distributions of force recruitment threshold 𝐹^𝑡ℎ^measured for the MUs identified in that study, in the tibialis Anterior (TA) and Vastus Lateralis (VL) muscles. Confirming previous thoughts (Heckman & Enoka, 2012; Duchateau & Enoka, 2022; Caillet et al., 2022a), the 𝐹^𝑡ℎ^threshold properties in Figure 2 are exponentially distributed, which suggests many low-threshold MUs and few high-threshold MUs in the MU pool, as half of the MU population is recruited below 25% MVC in both the human TA and VL muscles (Enoka & Fuglevand, 2000). A similar exponential distribution was observed for the current recruitment thresholds 𝐼_𝑡ℎ_ in Caillet et al. (2022a). It is worth mentioning that the experimental 𝐹^𝑡ℎ^and 𝐼_𝑡ℎ_ data points were slightly better fitted with a linear-power relationship than with an exponential function, although the terminology ‘exponential distribution of recruitment thresholds’ is maintained in the following for simplicity. In conclusion, human muscles have more low-threshold MUs than high-threshold MUs, with a continuous distribution of recruitment thresholds and in proportions that are subject- and muscle-specific.

**Figure 2:**
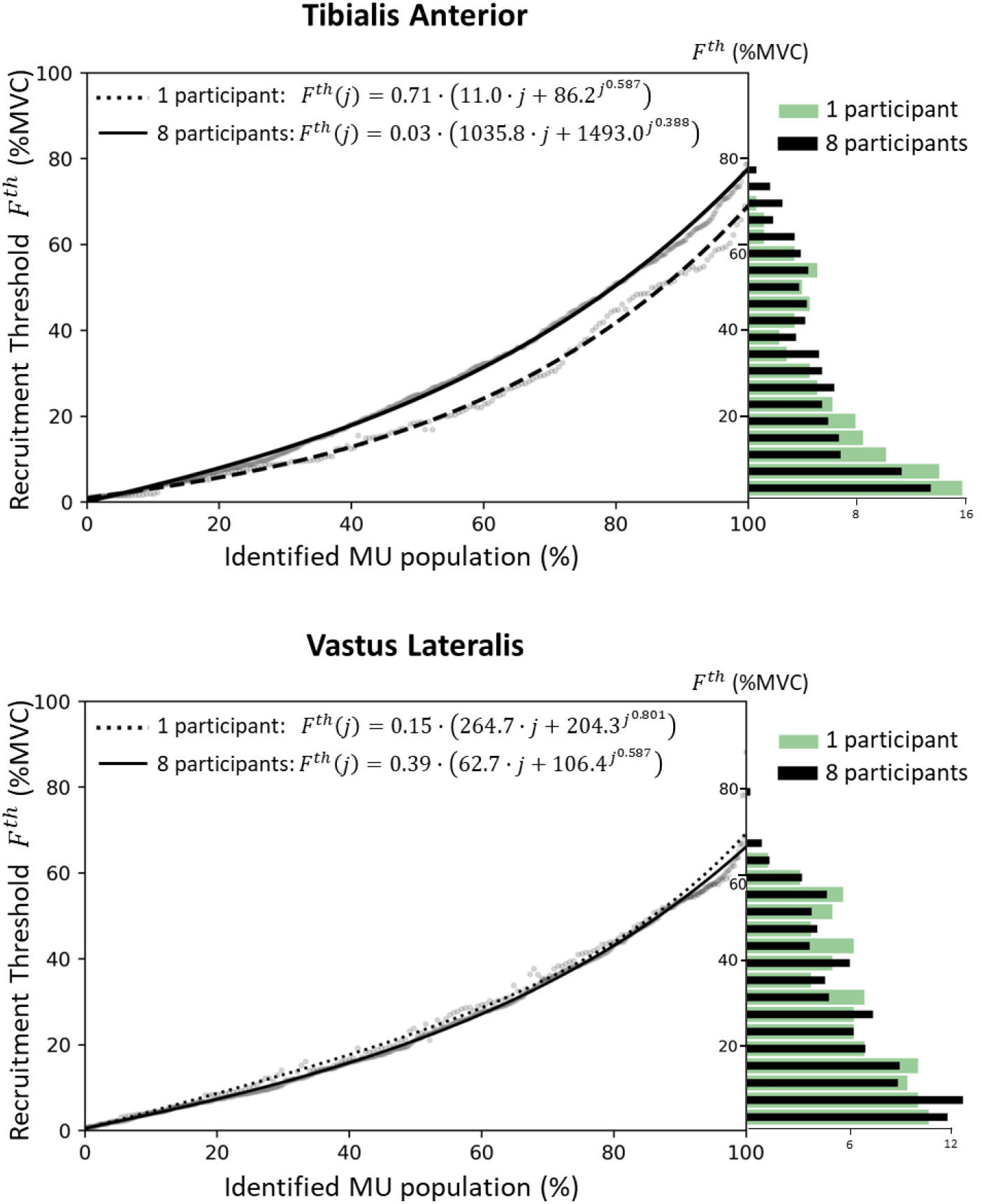
Continuous distribution of motor unit (MU) force recruitment thresholds 𝑭^𝒕𝒉^ across the MU pool of two human muscles. The experimental data was obtained by Avrillon et al. (2023a), where extensive samples of unique MUs (40-200 in a single participant, >800 for eight participants) were identified over a wide range of contraction intensities (10% to 80% MVC) for the tibialis anterior and vastus lateralis muscles. The density histograms report for a single participant (green bars) and for the cohort of eight participants (black bars) the percentage of identified MUs (x-axis) with respect to their measured force recruitment threshold 𝐹^𝑡ℎ^in percentage of the maximum voluntary contraction (%MVC) force (y-axis). Bins are 4% MVC-wide. An exponential distribution is observed; many MUs have low recruitment thresholds and few MUs have high recruitment thresholds. The scatter plots report for a single participant (light grey data points) and for the cohort of eight participants (black bars, dark grey data points) the relationship between the force recruitment threshold 𝐹^𝑡ℎ^ (in %MVC, y-axis) of a MU and the rank (in %, x-axis) it occupies in the identified MU pool, ranked in ascending order of recruitment thresholds. The clouds of data points were fitted with the linear-power relationships (r^2^>0.97, p<0.001) reported in the inset, where 𝑗 ∈ [0; 1]. Linear-power relationships returned better fits than purely exponential and linear-exponential functions. However, the terminology ‘exponential distribution of recruitment thresholds’ is used in this study for clarity. In a pool of 𝑁 MUs, these relationships can be adapted with the 𝑗 → ^𝑗^𝑁 transformation, where 𝑗 becomes an integer in ⟦1; 𝑁⟧. It is worth noting that joint torques were measured in Avrillon et al. (2023a); joint torques were assimilated to muscle forces, as the bipolar EMG signals concurrently recorded from the co-contracting agonist and antagonistic muscles did not show changes in relative activation with contraction intensities.

### Representative samples of MUs

As a practical definition, a MU sample is representative of the MU pool if the activity of the MUs of the sample retains most of the information contained in the activity of the full pool of MUs. This implies that a representative sample should be “sufficiently” large and with a distribution of recruitment thresholds similar to that of the full pool, i.e., continuous and approximately exponential. The first conditions on size and continuity are needed to avoid gaps in MU behaviour in specific recruitment force ranges. Because of the need to represent an exponential distribution, a sample of MUs is representative of the full MU pool if it contains many more low-threshold than high-threshold MUs. For example, the TA muscle comprises a total of ∼400 MUs. When it contracts at 50% of the maximum voluntary contraction (MVC) force, the active MUs would be approximately 320 (see Figure 2). In these conditions, a sample of 32 MUs (i.e., 10% of the active pool) would be representative of the MU pool if 10, 7, 6, 5, and 4 MUs were identified for each 10%-MVC interval of force, as given by the 𝐹^𝑡ℎ^- distribution of the full MU pool (solid curve) in Figure 2.

## Estimation of the neural drive to muscle

It is currently not possible to directly observe during a motor task all concurrently active MUs of a human muscle. Conversely, the neural drive to muscles can be indirectly approximated by electrophysiological features associated to its intensity, such as the amplitude of EMG signals, or from experimental samples of observed MU activities, or from computational estimates of the activities of non-observed MUs.

### Global analysis of EMG signals

The neural drive to muscles has been classically approximated from the myoelectric activity detected with electrodes applied on the skin covering the investigated muscle (surface EMG) or inserted into the muscle (intramuscular EMG) (Besomi et al., 2019; De Luca, 1997; Farina et al., 2004). In this context, recent tutorials (Merletti & Muceli, 2019; Merletti & Cerone, 2020; Clancy et al., 2023) and consensus studies (Besomi et al., 2020; Gallina et al., 2022) provide best practices for optimal choice of electrode design, montage, and placement, and for optimal EMG acquisition and post-processing. For example, the estimate of amplitude of the surface EMG, e.g., low-pass filtered rectified EMG, is commonly associated to the intensity of the neural drive to muscle (Farina et al., 2014b). However, EMG amplitude provides only a crude description of the neural drive. Indeed, surface EMG signals are influenced by the volume conductor properties as well as by amplitude cancellation, i.e., the amplitude of the interference EMG signal is less than the sum of the amplitudes of the MUAP trains (Farina et al., 2014b). The amount of amplitude cancellation differs between MUs and is greatest for small surface-recorded action potentials, which are associated with low-threshold small MUs and MUs located far from the recording electrodes. Consequently, amplitude cancellation reduces the contribution of some MUs to the estimate of EMG amplitude and the magnitude of the deficit cannot be predicted or compensated (Farina et al., 2014b). Also, the contribution of the discharging MUs to the EMG signal amplitude is determined by the amplitude of their MUAPs as filtered by the volume conductor and detected by the electrodes. Consequently, the large MUs that include many fibres and the MUs that are superficial and located close to the recording electrodes are over-represented in the recorded EMG signals. Conversely, surface EMG amplitude can be relatively insensitive to changes in the activity of small low-threshold MUs or MUs located far from the recording electrodes, i.e., deep MUs. For these reasons, the amplitude of surface EMG signals provides a relatively poor estimate of the intensity of the neural drive to muscle (Farina et al., 2014b). Estimates of the neural drive from the global analysis of EMG signals are sufficient in some applications but not in others, for example when building accurate subject-specific neuromusculoskeletal models of a motor task (Lloyd et al., 2023).

### Samples of identified MUs from decomposed high-density EMG signals

The combination of arrays of high-density EMG (HDEMG) electrodes with modern source-separation algorithms enables the identification of MU discharges in human *in vivo* (Holobar & Farina, 2014; Negro et al., 2016; Farina & Holobar, 2016). Contrary to the global analysis of EMG signals, samples of identified MUs provide a direct information on the discharge dynamics of the decoded MU population, which is a first step towards the identification of the neural drive to muscle. In brief, source-separation methods identify spike trains specific to individual MUs from the multi-unit EMG based on their characteristic MUAP shapes. Beside source separation methods, other approaches have been proposed to decompose the EMG signal, such as based on the detection of templates of MUAP waveforms (De Luca et al., 2006; Nawab et al., 2010). Consensus studies (Hug et al., 2021a; Gallina et al., 2022; Martinez-Valdes et al., 2023), tutorials (Del Vecchio et al., 2020; Avrillon et al., 2023b), open-source tools (Avrillon et al., 2023b; Valli et al., 2024) and commercial software packages (e.g., DEMUSE: https://demuse.feri.um.si/) for HDEMG acquisition, decomposition and/or spike train editing are available, with ongoing research towards fully automated methods for spike train identification (Clarke et al., 2020; Rossato et al., 2023).

### Classic limitations of EMG decomposition

Current source-separation algorithms tend to identify the MUs with the largest MUAPs within the EMG signal (Farina & Holobar, 2016). As high-threshold MUs have the largest innervation ratios in the MU pool (see Figure 2 in Caillet et al. (2023a)), decomposition techniques tend to favour the identification of the higher-threshold MUs. Therefore, by usually over-representing high-threshold over low-threshold MUs, the identified MU samples are likely not representative of the MU pool, as they do not follow an exponential distribution of recruitment thresholds (i.e., a greater proportion of low-threshold MUs over high-threshold MUs). In addition to this bias, it is usually difficult to identify large samples of MUs. With surface grids of 64 electrodes and/or intramuscular arrays of 40 electrodes, high-yield decomposition typically identifies samples of 5-40 MUs per muscle per contraction (Del Vecchio et al., 2020; Hug et al., 2021b; Muceli et al., 2022). This is still a limited portion of the MUs in human muscles that are typically hundreds to a few thousands. For these reasons, it is challenging to identify, with current EMG recording and decomposition techniques, MU samples that accurately estimate the neural drive to muscle (Figure 1, first column). For example, Caillet et al. (2022b) computed the neural drive at the muscle-scale with publicly available datasets of few identified MUs (i.e., 14-32 MUs), that over-represent higher-threshold with respect to lower-threshold MUs. The predicted neural drive, that was calculated as the low-pass filtered (10Hz) cumulative spike train and validated against the normalized recorded joint torque as previously proposed (Farina & Negro, 2015), substantially underestimated the true neural drive in the regions of low forces. Solutions to this problem are discussed in the following.

### Dense surface EMG grids of electrodes

Increasing the density and number of surface EMG electrodes yields larger and more representative samples of identified MUs (Figure 1, second column). Specifically, dense grids of surface EMG electrodes allow to better identify low-threshold MUs, while increasing the number of surface EMG electrodes allows to identify greater number of MUs. This was recently demonstrated by Caillet et al. (2023b), who showed decomposition of EMG signals recorded from the TA muscle with surface grids of 256 electrodes and 4-mm interelectrode distance (IED), as well as six grids of lower size, electrode density, and number of electrodes. The conclusions on electrode number and density were confirmed with computational simulations and with an ultra-dense grid of 2-mm IED in the same study (Caillet et al., 2023b), as well as for the VL muscle (unpublished results obtained from the publicly available data by Avrillon et al. (2023)). Figure 2 reports the density histograms (blue bars) of the MUs identified with three grids of 4-, 8-, and 12-mm IED and 256, 64, and 35 electrodes.

Increasing the electrode density gradually revealed early-recruited MUs that were not identified with lower-density grids. For example, up to 32% (range: 6-44%) of the MUs identified with the densest grid (256 electrodes, 4-mm IED) were early-recruited MUs (i.e., in the first half of the active MU pool), against only 11% (range: 0-24%) when using grids of 12-mm IED. Importantly, relatively more early-recruited MUs were identified than late-recruited MUs with increasing electrode densities. As a result, the densest grids of electrodes returned samples of identified MUs that spanned across the complete recruitment range, approaching the physiological exponential distribution of recruitment thresholds necessary for accurate neural drive estimation (see top-middle and top-right density histograms in Figure 3, blue bars). Conversely, the grid with 12-mm IED revealed few of the active low-threshold MUs recruited during the first half of the muscle contraction. Increasing the electrode density enables to better sample the action potential profiles of the small MUs and MUs far from the recording sites across multiple electrodes and increases the pulse-to-noise ratio of their identified spike trains, better separating them from the residual noise and enabling their identification (Caillet et al., 2023b).

**Figure 3:**
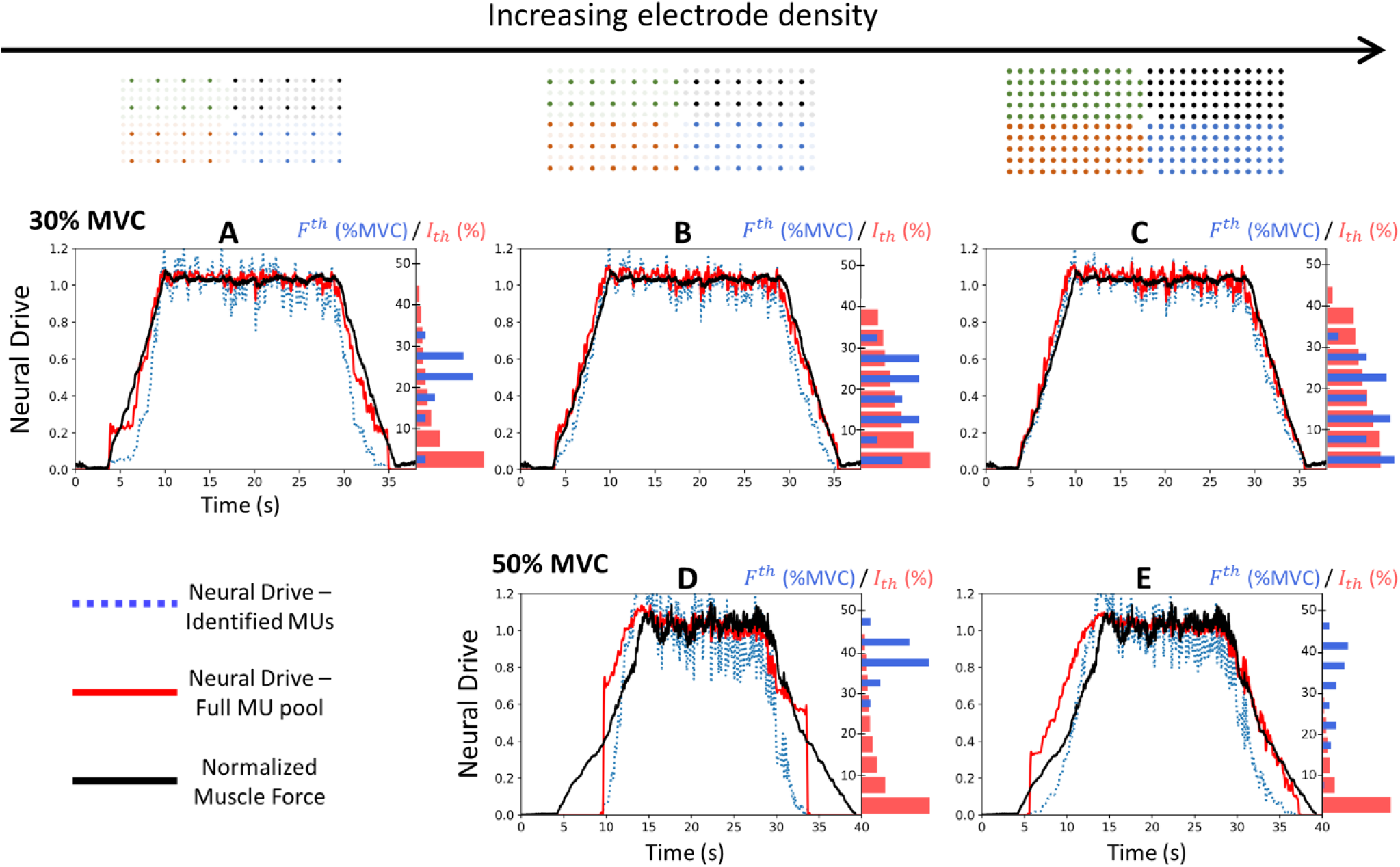
Estimation of the neural drive to muscle from populations of discharging motor units (MUs). The data and results were obtained in Caillet et al. (2023a, 2023b). EMG signals were recorded from the tibialis anterior muscle with three grid configurations of increasing electrode density (from left to right: grids of 12-mm, 8-mm, and 4-mm interelectrode distance involving 35, 64, and 256 electrodes, respectively) during trapezoidal contractions up to 30% maximum voluntary contraction (MVC) (first row) and 50% MVC (second row). Samples of MUs were identified from the decomposition of these EMG signals. The experimental neural drive was estimated from the samples of identified MUs (dotted blue traces) as the low-pass filtered cumulative spike train. The neural drive was also estimated from the discharge activity of the artificial full pool population of 𝑁 = 400 MUs (solid red traces), that was generated from the experimental MU samples with computational methods (Caillet et al., 2022b) (see text for details). The neural drive was validated in the normalized space against the measured TA force (Farina & Negro, 2015; Enoka & Farina, 2021) (solid dark trace, see text for details on the validation). The histograms report the density distribution of the MUs across the recruitment range in 5%-steps. The distribution of experimentally-identified MUs is reported with blue bars according to their measured force recruitment thresholds 𝐹^𝑡ℎ^ (in %MVC). The distribution of the artificial MUs from the reconstructed pool is reported with red bars according to their predicted current recruitment thresholds I_th_ (in percentage of the maximum theoretical value for 𝐼_𝑡ℎ_). The histograms were normalized to the total number of MUs.

Increasing the number of electrodes gradually revealed MUs that were not identified with fewer electrodes. Notably, the number of identified MUs doubled with 256 electrodes (4-mm IED and 36 cm^2^) against 64 electrodes (4-mm IED or 8-mm IED), reaching 56±14 MUs and 45±10 MUs for contractions up to 30% and 50% MVC, respectively. Besides, increasing the number of electrodes also helps identifying MUs for each interval of recruitment thresholds. It is worth noting that the rate at which additional MUs are detected with higher number of electrodes decreases when the grids become very dense (< 4-mm IED).

Estimations of the neural drive at the muscle scale improved with increasing the density of surface electrode grids and the number of electrodes (as displayed in Figure 2 and demonstrated by Caillet et al. (2023a)). For example, the low-density grid of 35 electrodes returned an inaccurate estimate of the neural drive (top-left subplot in Figure 3) with sudden jumps and drops over the interval of recruitment threshold where no MUs were identified, and underestimation in the regions of low forces where the low-threshold active MUs were not identified. By identifying more representative MU samples, more accurate estimates of the neural drive were obtained with the dense grid of 256 electrodes (top-right subplot in Figure 3).

### Dense intramuscular EMG arrays of electrodes

High-density arrays of intramuscular electrodes based on thin-film technology have also been proposed (Muceli et al., 2015, 2022; Chung et al., 2023). Like surface EMG, the number of identified MUs increases with the number of intramuscular recording sites. Negro et al. (2016) automatically decoded 20±3 (up to 24) MUs in the TA muscle from 16 to 32 intramuscular recording sites at forces ranging from 10 to 30% MVC, while Muceli et al. (2022) automatically decoded 31±5 (up to 40) MUs when using 40 channels at 30% MVC, and up to 67 MUs with 80 channels. Source-separation algorithms might detect relatively more low-threshold MUs from intramuscular EMG than surface EMG, as intramuscular EMG is not subject to the skin filtering and attenuating effects and has higher signal-to-noise ratio than surface EMG (Besomi et al., 2019). It remains to be investigated whether the MU samples identified with intramuscular high-density arrays of electrodes are representative of the full MU pool. Importantly, intramuscular arrays of electrodes can identify the activity of the deep MUs that cannot be detected from surface EMG. Measurements may therefore combine surface and intramuscular arrays of electrodes to detect separate MU populations in difference portions of the muscle (Muceli et al., 2015). Therefore, concurrently recording EMG signals from dense and large surface grids of electrodes and multiple intramuscular high-density arrays of electrodes targeting different populations of deep MUs can increase the experimental yield of MUs and the identification of a strongly representative sample of the MU pool (Negro et al., 2016; Muceli et al., 2022).

### Selecting optimal subsets of the identified MUs

While with modern approaches it is possible to increase the number of experimentally identified MUs and favour to some extent the identification of low-threshold MUs, as discussed above, the distributions of these experimental samples are rarely matching the distribution of MUs in the pool. For example, a TA muscle contracting at 30% MVC has a third of its active MUs recruited below 8% MVC, another third between 8 and 18% MVC, and another third between 18 and 30%MVC (Figure 2), i.e., a 33%-33%-34% split of the active MUs over these three recruitment ranges. Conversely, the 64- electrode grid returned, for the group of participants in Caillet et al. (2023b), a 10±5% - 26±10% - 64±10% split of identified MUs over those three recruitment ranges. This indicates that the higher-threshold MUs in the 18-30% MVC range are over-represented. As expected, the 256-electrode grid returned a more balanced 16±9% - 26±8% - 58±16% split, although the higher-threshold MUs were still over-represented (Caillet et al. 2023b). To address this limitation, it is possible to select an optimal MU subset of the identified MU sample, that is more representative of the MU pool, by discarding some of the identified MUs that fall into the recruitment threshold ranges that are over-represented. The selected optimal subsets would better estimate the neural drive to muscle than the original samples of identified MUs. This approach by optimal subsets proved beneficial in dexterous human-machine interfacing (Yeung et al., 2023), for example.

To illustrate this approach, we analysed here 500,000 random combinations of subsets of the MU samples identified in Caillet et al. (2023b), without repetitions within and between subsets, and with more than 15 MUs. The 200 combinations of MUs that best estimated the neural drive at muscle-scale (according to a weighted combination of normalized coefficient of determination and root-mean square error) were extracted. On average, the optimal subsets of MUs followed a 25±2% - 29±3% - 46±3% distribution over the three defined recruitment ranges, when the original MU sample was obtained with the 256-electrode grid. A similar 23±3% - 30±2% - 49±4% split was achieved by the optimal subsets of the 64-electrode grid. In one participant, for whom many low-threshold MUs were identified (top-right quadrant in Figure 3), the optimal subset achieved a close-to-theoretical 30%-32%- 38% split. Those optimal subsets better estimated the neural drive to muscle than the original MU samples by effectively discarding a portion of the over-represented higher-threshold MUs and retaining the identified lower-threshold MUs. This method by optimal subsets however reduces the available experimental data and impoverishes our direct experimental observation and understanding of the dynamics of neural control strategy in humans *in vivo*. The discarded experimental data may rather be used to infer the activity of the non-identified active MUs, as we will discuss later.

### Limitations of state-of-the-art recording and decoding methods

Despite the recent improvement discussed above, experimental approaches for the identification of representative samples of MUs still present several limitations. First, experimental MU decoding is mainly limited to isometric and static contractions. New computational approaches are emerging to identify individual MUs in quasi-static (Oliveira & Negro, 2021) and dynamic (Glaser & Holobar, 2018) tasks but other force trajectories more representative of daily-life contractions should be investigated. Second, the identification of many low-threshold MUs is not always possible. For example, very few low-threshold MUs were identified in Caillet et al. (2023b) during high-force contractions, even with the densest grids of electrodes (similar results are obtained at high forces by the intramuscular dense arrays). Consequently, inaccurate estimations of the neural drive were obtained for high-force contractions, irrespective of the electrode density, in the regions of low forces where very few low-threshold MUs were identified (Figure 3, bottom row). The percentage of identified low-threshold MUs is also highly subject-specific, with a high disparity of results reported between subjects (Caillet et al., 2023b). It is likely muscle-specific, although no significant difference was found between the TA and VL muscles (Avrillon et al., 2023a), and gender-specific, as fewer low-threshold MUs are usually identified in females than males using non-invasive methods (Jenz et al., 2023). To ensure the identification of many MUs over the full recruitment range, one can track identified MUs across multiple contraction intensities. For example, Avrillon et al. (2023a) applied this method with the 256- electrode montage over the TA (VL) muscle of 16 participants for ten contractions between 10% MVC and 80% MVC, and identified samples of 129±44 (130±63) unique MUs per subject and muscle, the density distribution of which displayed the physiological exponential tendency (black bars in Figure 2) typical of the full MU pool.

## Computational augmentation of experimental MU samples

Although representative samples of MUs can be experimentally identified in some conditions, it remains impossible to experimentally detect the activity of full pools of active MUs, i.e., to experimentally identify the neural drive at the MU scale. To address this limitation, computational methods of data augmentation (Caillet et al., 2022b; Ornelas-Kobayashi et al., 2023) have been recently developed to estimate the discharge activity of the full MU pool from a limited sample of identified MUs (Figure 1, third column). An open-source implementation of the approach proposed by Caillet et al. (2022b) is publicly available at https://github.com/ArnaultCAILLET/Caillet-et-al-2022-PLOS_Comput_Biol.

### Methods of experimental data augmentation

Caillet et al. (2022b) proposed a computational data-driven method that uses the physiological information of a sample of identified MUs to computationally estimate realistic discharge activities (i.e., spike trains) for the remaining non-identified MUs. This method is briefly described in the following and is applied in Figure 4 with an input sample of 81 MUs identified from surface HDEMG signals (Figure 4A).

**Figure 4:**
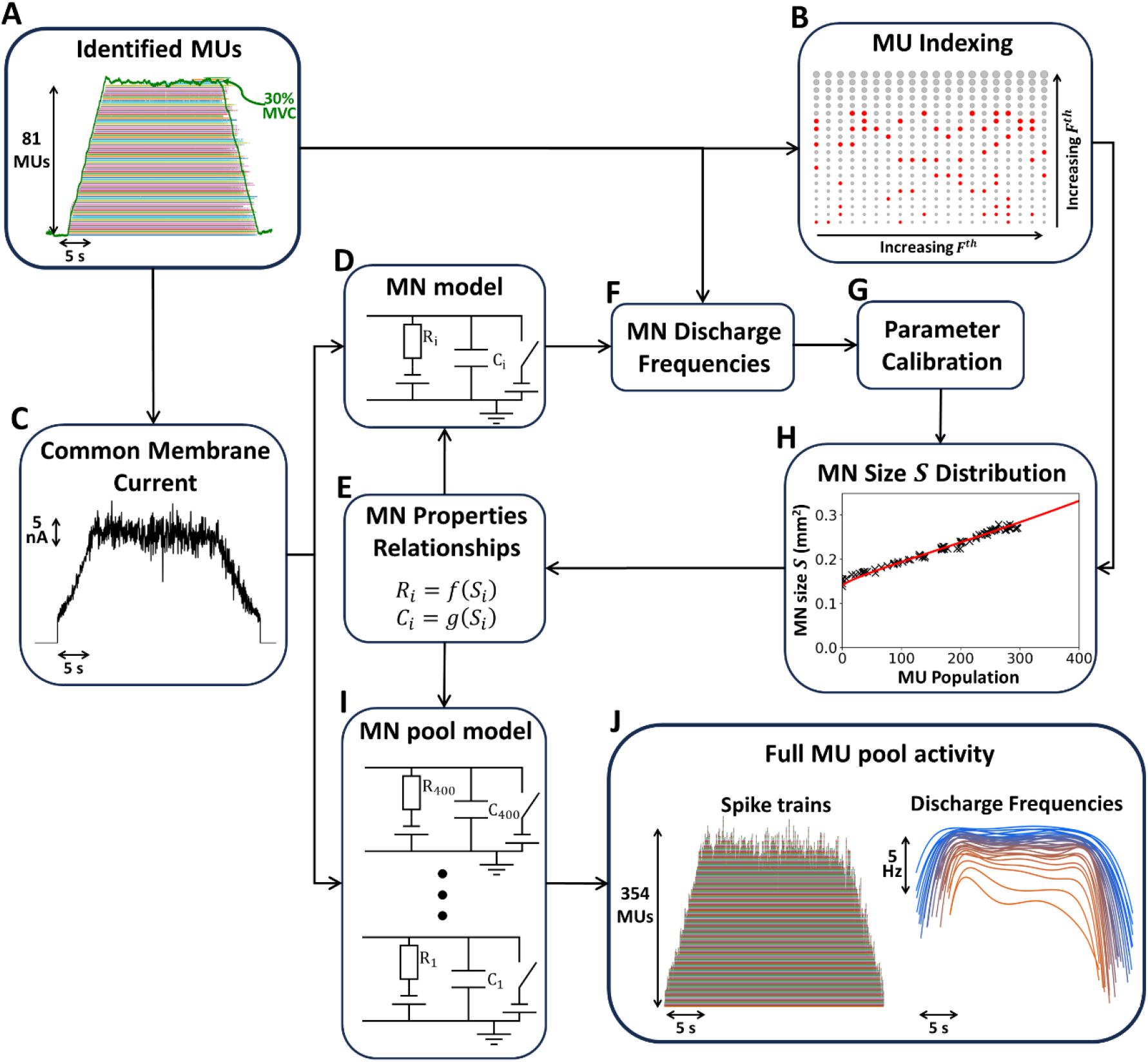
Computational method for reconstructing the discharge activity of the full pool of motor units (MUs) from a limited sample of experimentally-identified MUs. (Caillet et al., 2023a). (A) In this example, the spike trains of 81 MUs were identified from the decomposition of high-density EMG signals (256 electrodes, 4-mm interelectrode distance, 36 cm^2^, see Figure 3) recorded during a trapezoidal isometric contraction of the tibialis anterior muscle up to 30% maximum voluntary contraction (MVC). (B) The 81 identified MUs (red circles) are mapped with an index 𝑖 ∈ ⟦1; 𝑁⟧ into the MU pool of 𝑁 = 400 MUs according to their measured force recruitment threshold 𝐹^𝑡ℎ^. The grey circles denote MUs that were not identified experimentally. (C) The MN membrane current, common to all modelled MNs, is derived from the Common Synaptic Input to the MU pool, which is inferred from the discharge activity of the 81 identified MUs. (D) The Common Synaptic Input drives a revisited leaky-integrate-and-fire model of motoneuron (MN), the electrophysiological properties (e.g., input resistance 𝑅, membrane capacitance 𝐶) of which are all mathematically related to the size of the MN 𝑆, according to a framework of mathematical relationships between inter-related MN-specific properties derived elsewhere (Caillet et al., 2022a) and discussed in Section 2.1. (F) The difference between model-predicted and experimental MN discharge frequencies is minimized by (G) calibrating the size parameter 𝑆 for the 81 identified MUs. (H) The discrete 𝑆- distribution is fitted, resulting in a continuous distribution for the 𝑆 parameter. The resulting continuous distribution of MN-specific electrophysiological properties (E) scales a pool of MN models (I), that transforms the Common Synaptic Input (C) into the discharge activity of the full MU pool (J). It is worth noting that the method presented here, which considers a single common input driving a group of MNs, may be extended to multiple groups of MNs (e.g., MN clusters in a MN pool (Hug et al., 2023)), modelled as separate cohorts of MN LIF models and receiving specific common membrane currents that would be estimated separately.

The experimentally identified MUs are first indexed (𝑖 ∈ ⟦1; 𝑁⟧) in the full pool of 𝑁 MUs, according to their recorded recruitment threshold value 𝐹^𝑡ℎ^ and a typical 𝐹^𝑡ℎ^-distribution across the MU pool (e.g., Figure 2). Then, the common drive responsible for the discharge activity of the identified MUs is estimated (Figure 4C). To do so, the neural drive is computed from the identified spike trains (low-pass filtered cumulative spike train) and linearly associated to the common synaptic input in arbitrary values (Farina & Negro, 2015). Then, a phenomenological ‘Common Membrane Current’ (in nano Ampers, Figure 4C) is piecewise-linearly scaled from the common synaptic input with physiological values of MN current recruitment threshold (Caillet et al., 2022a). The method determines, for each identified MU, the set of MN-specific electrophysiological properties that best explain its observed discharge activity. These properties (i.e., membrane time constant, input resistance, capacitance, current recruitment threshold) are the parameters of a single-compartment leaky-integrate-and-fire (LIF) model of the MN (Figure 4D). Those inter-related properties are entirely determined by the MN size parameter 𝑆, as direct result from the discussion in Section 2.1 about the mathematical inter-relation between MN properties (Figure 4E, (Caillet et al., 2022a)). The calibrated LIF model transforms the Common Membrane Current (Figure 4C) into a train of discharge events. After calibration of the size parameter 𝑆 (Figure 4G), the MN-specific LIF model minimizes the root-mean-square difference between experimental and LIF-predicted filtered discharge frequencies (Figure 4F). Next, the scatted pairs {𝑖; 𝑆_𝑖_} (black crosses in Figure 4H) of MN index 𝑖 (Figure 4B) and calibrated parameter 𝑆_𝑖_ (Figure

4D-G) are fitted with a continuous function (red solid trace in Figure 4H). The resulting 𝑆-distribution, to which the previously-introduced mathematical relationships are re-applied (Figure 4E), determines the continuous distributions across the MU pool’s recruitment range of the MN electrophysiological parameters of the LIF model. Finally, the discharge activity of the full pool of active MUs is predicted. A population of 𝑁 LIF models of MNs (Figure 4I) is scaled with the continuous distributions of MN-specific electrophysiological properties derived in the last step. The digital twin of the full pool of 𝑁 MUs transforms the Common Membrane Current into a physiologically plausible estimation of the discharge activity of the full pool of active MUs (Figure 4J).

Ornelas-Kobayashi et al. (2023) proposed a computational method similar in some steps to that of Caillet et al. (2022b), applied to a 15-parameter single-compartment conductance-based model with four ion-gating channel types (Cisi & Kohn, 2008). This model of MN is more physiologically accurate than the LIF model shown in Figure 4D, however it is substantially more complex. As ion-gating channel-related parameters are difficult to measure in mammals (Bondarenko et al., 2004; Fohlmeister, 2009), Ornelas-Kobayashi et al. (2023) kept 12 MN parameters generic and constant, as in the original model (Cisi & Kohn, 2008), and identified the three remaining parameters with separate calibration routines. Consequently, the sets of MN parameters were not inter-related and were only partially MN-specific in that study, contrary to those derived in Figure 4. Conversely, when MN-specific parameters are inter-related (like in Figure 4E), a single calibration procedure is sufficient for accurate MN-specific predictions.

In conclusion, depending on the objective of the study, a trade-off must be found on MN model complexity. When reconstructing the full MU pool for estimating the neural drive to muscle, reconstructing the comprehensive exponential distribution of the MU recruitment thresholds is of critical importance, as achieved in Caillet et al. (2022b) (red bar histograms in Figure 3). Conversely, in Ornelas-Kobayashi et al. (2023), the trapezoidal contractions of very short ramps, where all active MUs are recruited over short 0.5-to-2-second periods, prevent from investigating the recruitment activity of the MU pool and are unfit to assess the distribution of the reconstructed full pool of MUs across the recruitment range. In many applications, detailing the physiological mechanisms responsible for firing action potentials is not necessary, as long as the predicted spike trains are accurate. In such cases, the flexibility of simple LIF models for receiving distributions of MN-specific inter-related MN parameters (Figure 4D,E) is more suitable for accurately reconstructing the MU pool activity, while additional MN modelling complexity is not warranted and may become a limiting factor due to the increased number of independent parameters to identify via optimization approaches.

The two reconstruction methods described above are currently the only methods in the literature that drive a computational model of MN pool with motoneuronal data that is obtained experimentally. MN pool models of varying complexity were previously proposed (Fuglevand et al., 1993; Cisi & Kohn, 2008; Dideriksen et al., 2010; Dideriksen et al., 2011; Negro & Farina, 2011; Watanabe et al., 2013; Farina et al., 2014a; Potvin & Fuglevand, 2017), although they were tested and driven with artificial inputs, inputs from interneurons, or feedback systems.

### Estimation of the neural drive to muscle

The full pool of artificial MUs obtained with the reconstruction method summarised in Figure 4 provides a more accurate estimate of the neural drive at the muscle-scale than the original experimental sample of identified MUs (red solid traces versus blue dotted traces in Figure 3). This was demonstrated with 15 MU samples identified from EMG signals recorded from the TA muscle with grids of varying electrode densities in three subjects performing trapezoidal contraction forces at up to 30% and 50% MVC (Caillet, 2023). The higher accuracy in estimating the neural drive at muscle-scale is partially explained by the exponential distribution of recruitment thresholds obtained for the full pool of 𝑁 artificial MUs (histograms of Figure 3, red bars). Of note, the predicted quasi-linear 𝑆- distribution in Figure 4H nonlinearly transforms into an exponential distribution of current recruitment thresholds 𝐼_𝑡ℎ_ with the mathematical relationship 𝐼_𝑡ℎ_ ∝ 𝑆^2.52^ (Caillet et al., 2022a).

The full pool of artificial MUs provides a comprehensive description of the neural drive at the MU-scale, by estimating the discharge activity for each active MU of the pool. Importantly, these estimates of discharge activity are accurate; this was demonstrated with a leave-one-out cross-validation that confirmed the accuracy of the estimated discharge activity for each of the identified MUs (Caillet et al., 2022b). This physiological accuracy is achieved because the method relies on physiologically-realistic mechanisms. Namely, the framework of mathematical relationships between inter-related MN-specific properties (Figure 4E) maintains the parameters and predictions of the LIF model within physiological bounds defined by the available experimental data. Moreover, this mathematical framework ensures a physiological distribution of the MN properties across the MU pool, consistent with the Henneman’s size principle (Henneman, 1981). Consequently, the method outputs MU pool discharge activities that are consistent with physiological laws (Figure 4J). For these reasons, reconstruction methods, such as that described in Figure 4, are powerful tools for accurately describing the neural drive to muscle both at the MU-scale and the muscle-scale from limited samples of experimentally observed MUs.

### Future perspectives of MU pool reconstruction methods

Methods of experimental data augmentation for MU analysis have potential for further developments. First, the accuracy of the neural drive estimated from the artificial full pool of MUs is sensitive to the quality of the observed sample of identified MUs. In Figure 3, the neural drive is accurately predicted by the full MU pool as long as some MUs are experimentally identified across the full recruitment range, reaching high accuracy when many low-threshold MUs are identified (Figure 3, top-middle and top-right subplots), and good accuracy even when some low-threshold MUs are identified but largely under-represented (Figure 3, top-left subplot). However, the neural drive predicted by the full pool of artificial MUs remains inaccurate in the regions of low forces where the number of detected MUs is insufficient (bottom plots in Figure 3). In such cases, the reconstructed MU pools display exponential MU distributions, contrary to the input samples of experimentally identified MUs, but receive non-physiological Common Current Input (Figure 4C), that onset at the time of the earliest-recruited identified MU and is zero until that time. Second, the single-compartment LIF model with current synapses proposed in Figure 4C provides accurate predictions of MN firing behaviour but comes with intrinsic limitations in physiological accuracy (Burkitt, 2006; Teeter et al., 2018; Caillet et al., 2022b). For example, the saturation of firing rates is described phenomenologically in the LIF model of Figure 3, while it could be included in a single-compartment LIF model with lumped active conductances (Burkitt, 2006; Caillet et al., 2022b), where this nonlinearity can be described by decreasing the intensity of the input current with increasing membrane depolarization. Finally, it is relevant to note that the concepts and methods presented in this section and in Figure 4 relates to a single pool of MN that receives one source of common synaptic input. However, the same approaches can be extended in a straightforward way to multiple groups/pools of motor neurons, each receiving different sources of common input (Hug et al., 2023). In this case, once the separate groups of MNs are identified as each receiving common input, the MNs would be modelled by separate cohorts of MN LIF models.

## Neuromechanics: Predicting forces from accurate decoding of the neural drive

We reviewed experimental and computational methods to observe the behaviour of discharging MUs that provide accurate estimates of the neural drive to muscle during human muscle contractions. These identification methods are of particular importance for studies that aim to decode muscle function from neural intent, i.e. neuromechanical investigations on how the neural commands determine behaviour.

### Single-input single-output EMG-driven predictions of force

Traditionally, computational simulations of human muscle contractions are performed with EMG-driven Hill-type models of whole muscles (Caillet et al., 2022c). In these methods, a single-actuator Hill-type muscle model phenomenologically transforms, in a single-input single-output strategy, the experimental EMG envelope into muscle force. This approach is well-established (Caillet et al., 2022c), and has had important applications, for example in human-machine interfacing (Sartori et al., 2012; Ao et al., 2017; Caggiano et al., 2022; Cimolato et al., 2022). However, as discussed above, EMG envelopes provide only a crude description of the neural drive to muscle, so that force predictions and neuromechanical analyses are limited with this approach. As an example, because of lack of accuracy, it is not possible with this approach to analyse the effect of MU synchronization on the steadiness of a task (Thompson et al., 2018).

To address the limitations of classic EMG-driven models, some studies (Sartori et al., 2017; Thompson et al., 2018; Kapelner et al., 2020) replaced the EMG envelope with a more accurate description of the neural drive, that is the filtered cumulative spike train (fCST) computed from discharging MUs identified from decomposed HDEMG signals. Yet, in both the global EMG- and fCST-driven approaches, the recruitment and discharge dynamics of the active MU pool are lumped into a single phenomenological macroscopic neural control signal. These single-input single-output strategies therefore maintain muscle modelling at the macroscopic level rather than describing the force-generating process at the level of individual MUs. This hinders the investigation of human neuromuscular control with computational tools.

### Motoneuron-driven models of muscle force

In response to the limitations of single-input strategies, Caillet et al. (2023a) developed a neuromuscular model that comprehensively describes the force-generating dynamics of individual MUs. This method is summarised in Figure 5. A cohort of Hill-type models transforms an input vector of MN spike trains (Figure 5D,E) derived from HDEMG (Figure 5A, B) into an output vector of individual MU forces (Figure 5F), that sum into the predicted whole muscle force (Figure 5J). Models of MU pools were previously proposed (Hatze, 1978; Hatze, 1980; Fuglevand et al., 1993; Raikova et al., 2018; Carriou et al., 2019), but all were used in combination with an artificial (not data-driven) neural drive. In Figure 5, the neuromuscular model receives processed HDEMG data and was tested both with MU samples identified from HDEMG (Figure 5B,D) and full pool of artificial MUs (Figure 5E) obtained with the reconstruction method presented in Figure 4. These approaches were tested (Caillet et al., 2023a) for trapezoidal contractions of the TA muscle up to 30% and 50% MVC with three grids of varying interelectrode distance (as in Figure 3). An open-source implementation of this approach is publicly available at https://github.com/ArnaultCAILLET/MN-driven-Neuromuscular-Model-with-motor-unit-resolution. Across all conditions, the accuracy in predicting muscle force (Figure 5H, I) systematically increased, especially in the regions of low forces, with greater numbers of electrodes, denser grids of electrodes, with the full pool of artificial MUs obtained by data augmentation (red trace in Figure 5J) versus the experimental MU sample (blue trace in Figure 5J), and at 30% MVC versus 50% MVC. These results echo with the conclusions previously drawn and stress the importance of experimentally sampling the MU pool in a representative way for controlling MN-driven neuromuscular models with realistic neural drive. In particular, neglecting the discharge activity of low-threshold MUs results in ignoring the force-generating contribution of those MUs to the whole muscle force, that is underpredicted at low contraction intensities. Limited samples of identified MUs (e.g., samples of 2- 22 MUs used in another model of MU pool (Gogeascoechea et al., 2023)), which likely over-represent the activity of high-threshold MUs, are therefore unfit for accurately controlling models of a MU pool directly. This is why in cases of limited MU samples, the unexplained EMG residual is added to the fCST after parameter calibration in fCST-driven approaches (Sartori et al., 2017). As a final note, MN-driven models with MU resolution provide a more physiologically accurate description of the force-generating dynamics of muscles than traditional single-input single-output EMG-driven models. The functioning of traditional EMG-driven models relies on phenomenological non-linear transformations, like between EMG and “muscle activation”, that include phenomenological parameters that cannot be experimentally estimated and must be calibrated to achieve prediction accuracy, usually by matching estimated and ground truth joint moments. Parameter calibration yields a risk of parameter over- fitting when the calibration set is small and lacks a large variety of motor tasks, so that EMG-driven models may not perform well on test sets that include different trials of motor task (Caillet et al., 2023c). Conversely, MN-driven models with MU resolution may integrate parameters that have a physiological meaning and may be measured experimentally. For example, Caillet et al. (2023a) proposed advanced physiological models of the MUs’ activation dynamics with parameters directly identified from recent experimental data and achieved accurate estimations of force without having to perform steps of parameter calibration.

**Figure 5:**
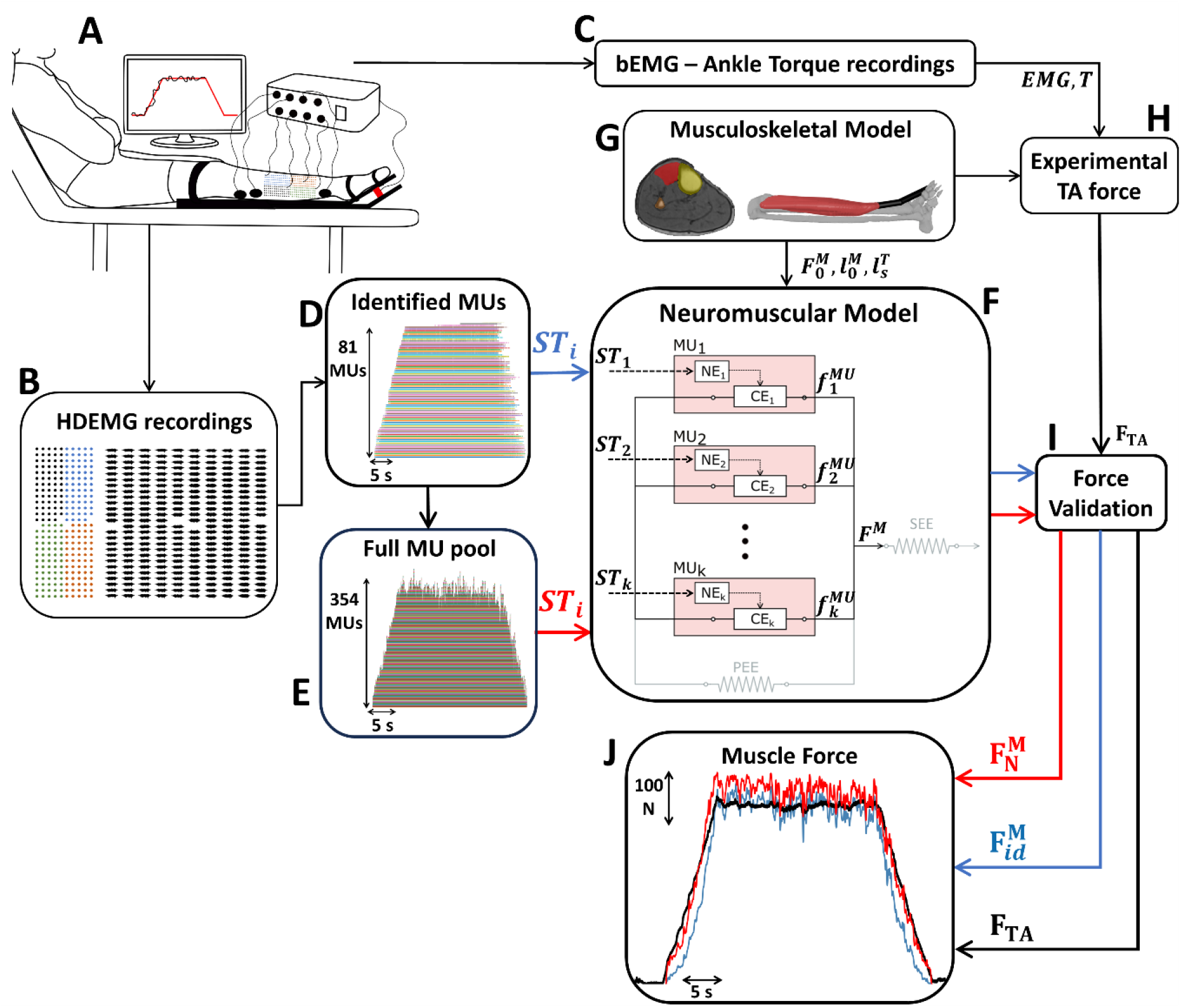
Computational method for predicting whole muscle forces from motoneuronal spiking activity using a motoneuron (MN)-driven computational muscle model with motor unit (MU) resolution. (A-C) In Caillet et al. (2023a), high-density electromyographic (HDEMG) signals were recorded from the tibialis anterior (TA) muscle during isometric trapezoidal dorsiflexion. The ankle torque 𝑇 and four bipolar EMG (bEMG) signals from co-contracting muscles were also recorded. (D) The HDEMG signals were decomposed into spike trains (𝑆𝑇_𝑖_, 𝑖 ∈ ⟦1; 81⟧) for 81 identified Mus in this example. (E) The discharge activity of the full pool of active MUs (𝑆𝑇_𝑖_, 𝑖 ∈ ⟦1; 354⟧) was generated from the 81 identified spike trains with the computational method in Figure 4. (F) A MN-driven neuromuscular model with MU resolution transforms the spike trains derived previously (𝑆𝑇_1_, 𝑆𝑇_2_, …, 𝑆𝑇_𝑘_) into MU forces (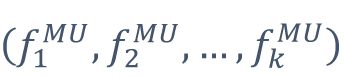), that sum into the predicted whole muscle force 𝐹^𝑀^. Here, 𝑘 = 81 or 𝑘 = 354 whether the neuromuscular model receives the spike trains from the experimental sample of identified MUs (blue arrow) or from the full pool of artificial MUs (red arrow). The neuromuscular model describes the activation and contraction dynamics of each modelled MU with dedicated Neural Element (NE) and Contractile Element (CE). The passive dynamics of the parallel and in-series elastic elements (PEE and SEE) are neglected in Caillet et al. (2023a). In this approach, the neuromuscular model receives a subject-specific MU-specific neural control. (G) A subject-specific musculoskeletal model, derived from segmented medical images, provides further subject-specific muscle-specific architectural parameters (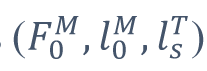) to couple with the neuromuscular model. (H) The subject-specific musculoskeletal model, the recorded ankle, and the bEMG signals recorded from co-contracting muscles are used to estimate the experimental force 𝐹_𝑇𝐴_ generated by the TA muscle (Caillet et al., 2023a). (I-J) The muscle force predicted from the experimental sample of identified MUs (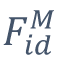, blue arrow and trace) and from the full pool of artificial MUs (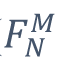, red arrow and trace) are validated against the experimental muscle force (𝐹_𝑇𝐴_, black arrow and trace).

## Conclusions and perspectives

The neural drive to muscle refers to the discharge activity of the active motoneuron pool, i.e., to the ensemble of action potentials transmitted to the muscle fibres and responsible for muscle force generation. The neural drive to muscles is the ultimate signal for muscle control; its accurate estimate is necessary for establishing its physiological link with the resulting muscle forces for the neuromechanical investigation of movement. Accurately identifying the neural drive to muscle in humans *in vivo* is however challenging. The MU samples identified from decomposed EMG signals usually include relatively few identified MUs, lack the activity of low-threshold MUs, and over- represent high-threshold MUs, so that they are inaccurate descriptions of the active MU pool. Here, we reviewed electrophysiological methods to identify larger samples of MUs that follow a physiological exponential distribution across the MU pool’s recruitment range, i.e., many low-threshold and few high-threshold MUs (Figure 2). These improvements are obtained by increasing the number and density of surface or intramuscular EMG electrodes (Figure 3). Then, we described how the discharge activity of the full MU pool can be reconstructed with computational data augmentation methods from the optimized – but still limited – experimental data (Figure 4).

The accurate and comprehensive description of the neural drive to muscle from individual motoneuronal activity finds crucial applications in the control of neuromuscular models (Figure 5), but also in the investigation of neural control strategies, including the study of neural synergies (Hug et al., 2023), functional connectivity (Bräcklein et al., 2022), or in neurorehabilitation technologies, e.g., for tremor suppression (Puttaraksa et al., 2022), prosthetics control (Farina et al., 2017; Farina et al., 2021), and assistive exoskeleton control (Jung et al., 2021).

## References

1. Ao D, Song R, Gao J (2017) Movement performance of human-robot cooperation control based on EMG- driven hill-type and proportional models for an ankle power-assist exoskeleton robot. TNSRE 25:1125–1134.

2. Avrillon S, Hug, Enoka, Caillet, Farina (2023a) The decoding of extensive samples of motor units in human muscles reveals the rate coding of whole motoneuron pools. Biorxiv .

3. Avrillon S, Hug F, Gibbs C, Farina D (2023b) Tutorial on MUedit: An open-source software for identifying and analysing the discharge timing of motor units from electromyographic signals. bioRxiv 2023.07. 13.548568.

4. Besomi M, Hodges PW, Clancy EA, Van Dieën J, Hug F, Lowery M, Merletti R, Søgaard K, Wrigley T, Besier T (2020) Consensus for experimental design in electromyography (CEDE) project: Amplitude normalization matrix. Journal of Electromyography and Kinesiology 53:102438.

5. Besomi M, Hodges PW, Van Dieën J, Carson RG, Clancy EA, Disselhorst-Klug C, Holobar A, Hug F, Kiernan MC, Lowery M (2019) Consensus for experimental design in electromyography (CEDE) project: Electrode selection matrix. Journal of Electromyography and Kinesiology 48:128–144.

6. Binder MD, Powers RK, Heckman CJ (2020) Nonlinear input-output functions of motoneurons. Physiology 35:31–39.

7. Binder, M. D., Heckman, C. J. & Powers, R. K. (1996) The physiological control of motoneuron activity. In Rowell LB, S. J. (ed), Handbook of Physiology: Exercise, Regulation and Integration of Multiple Systems. New York: Oxford Univ Press, pp. 3–53.

8. Bräcklein M, Barsakcioglu DY, Del Vecchio A, Ibáñez J, Farina D (2022) Reading and modulating cortical β bursts from motor unit spiking activity. Journal of Neuroscience 42:3611–3621.

9. Burke, R. E. (1981) Motor units: Anatomy, physiology, and functional organization. In Brooks, V. B. (ed), Handbook of Physiology, the Nervous System, Motor Control. American Physiological Society, pp. 345–422.

10. Burkitt AN (2006) A review of the integrate-and-fire neuron model: I. homogeneous synaptic input. Biol.Cybern. 95:1–19.

11. Caggiano V, Wang H, Durandau G, Sartori M, Kumar V (2022) MyoSuite--A contact-rich simulation suite for musculoskeletal motor control. arXiv preprint arXiv:2205.13600.

12. Caillet Arnault Hubert (2023) Neuromuscular modelling of skeletal muscle contraction from experimental motoneuronal activity. PhD Thesis, Imperial College London, Chapter 3.

13. Caillet AH, Phillips AT, Farina D, Modenese L (2022b) Estimation of the firing behaviour of a complete motoneuron pool by combining electromyography signal decomposition and realistic motoneuron modelling. PLOS Computational Biology 18:e1010556.

14. Caillet AH, Avrillon S, Kundu A, Yu T, Andrew T.M. Phillips, Modenese L, Farina D (2023b) Larger and denser: An optimal design for surface grids of EMG electrodes to identify greater and more representative samples of motor units. eNeuro ENEURO.0064–23.2023.

15. Caillet AH, Phillips ATM, Farina D, Modenese L (2023a) Motoneuron-driven computational muscle modelling with motor unit resolution and subject-specific musculoskeletal anatomy. PLOS Computational Biology 19:e1011606.

16. Caillet AH, Phillips AT, Farina D, Modenese L (2022a) Mathematical relationships between spinal motoneuron properties. eLife 11:e76489.

17. Caillet AH, Phillips AT, Carty C, Farina D, Modenese L (2022c) Hill-type computational models of muscle- tendon actuators: A systematic review. bioRxiv.

18. Carriou V, Boudaoud S, Laforet J, Mendes A, Canon F, Guiraud D (2019) Multiscale hill-type modeling of the mechanical muscle behavior driven by the neural drive in isometric conditions. Computers in biology and medicine 115:103480.

19. Chung, B., Zia, M., Thomas, K. A., Michaels, J. A., Jacob, A., Pack, A., … & Sober, S.J. (2023). Myomatrix arrays for high-definition muscle recording. Elife, 12, RP88551.

20. Cimolato A, Driessen JJ, Mattos LS, De Momi E, Laffranchi M, De Michieli L (2022) EMG-driven control in lower limb prostheses: A topic-based systematic review. Journal of NeuroEngineering and Rehabilitation 19:1–26.

21. Cisi RRL, Kohn AF (2008) Simulation system of spinal cord motor nuclei and associated nerves and muscles, in a web-based architecture. J Comput Neurosci 25:520–542.

22. Clancy EA, Morin EL, Hajian G, Merletti R (2023) Tutorial. surface electromyogram (sEMG) amplitude estimation: Best practices. Journal of Electromyography and Kinesiology 72:102807.

23. Clarke AK, Atashzar SF, Del Vecchio A, Barsakcioglu D, Muceli S, Bentley P, Urh F, Holobar A, Farina D (2020) Deep learning for robust decomposition of high-density surface EMG signals. IEEE Transactions on Biomedical Engineering 68:526–534.

24. De Luca CJ, Hostage EC (2010) Relationship between firing rate and recruitment threshold of motoneurons in voluntary isometric contractions. J.Neurophysiol. 104:1034–1046.

25. De Luca CJ (1997) The use of surface electromyography in biomechanics. Journal of applied biomechanics 13:135–163.

26. De Luca, Adam, Wotiz, Gilmore, Nawab, Decomposition of surface EMG signals. J Neurophysiol 96, 1646–1657 (2006).

27. Del Vecchio A, Holobar A, Falla D, Felici F, Enoka RM, Farina D (2020) Tutorial: Analysis of motor unit discharge characteristics from high-density surface EMG signals. Journal of Electromyography and Kinesiology 53:102426.

28. Del Vecchio A, Negro F, Felici F, Farina D (2018) Distribution of muscle fibre conduction velocity for representative samples of motor units in the full recruitment range of the tibialis anterior muscle. Acta Physiologica 222:e12930.

29. Dideriksen JL, Enoka RM, Farina D (2011) Neuromuscular adjustments that constrain submaximal EMG amplitude at task failure of sustained isometric contractions. Journal of Applied Physiology 111:485–494.

30. Dideriksen JL, Farina D, Baekgaard M, Enoka RM (2010) An integrative model of motor unit activity during sustained submaximal contractions. J.Appl.Physiol. 108:1550–1562.

31. Duchateau J, Enoka RM (2022) Distribution of motor unit properties across human muscles. J.Appl.Physiol. 132:1–13.

32. Enoka RM, Farina D (2021) Force steadiness: From motor units to voluntary actions. Physiology 36:114–130.

33. Enoka RM, Duchateau J (2017) Rate coding and the control of muscle force. Cold Spring Harbor perspectives in medicine 7:.

34. Enoka, R. M., & Fuglevand, A. J. (2001). Motor unit physiology: some unresolved issues. Muscle & Nerve: Official Journal of the American Association of Electrodiagnostic Medicine, 24(1), 4–17.

35. Farina D, Vujaklija I, Brånemark R, Bull AMJ, Dietl H, Graimann B, Hargrove LJ, Hoffmann K, Huang H, Ingvarsson T, Janusson HB, Kristjánsson K, Kuiken T, Micera S, Stieglitz T, Sturma A, Tyler D, Weir RFf, Aszmann OC (2021) Toward higher-performance bionic limbs for wider clinical use. Nature biomedical engineering 1–13.

36. Farina D, Vujaklija I, Sartori M, Kapelner T, Negro F, Jiang N, Bergmeister K, Andalib A, Principe J, Aszmann OC (2017) Man/machine interface based on the discharge timings of spinal motor neurons after targeted muscle reinnervation. Nature biomedical engineering 1:1–12.

37. Farina D, Holobar A (2016) Characterization of human motor units from surface EMG decomposition. Proc IEEE 104:353–373.

38. Farina D, Negro F (2015) Common synaptic input to motor neurons, motor unit synchronization, and force control. Exerc.Sport Sci.Rev. 43:23–33.

39. Farina D, Merletti R, Enoka RM (2014b) The extraction of neural strategies from the surface EMG: An update. J.Appl.Physiol. 117:1215–1230.

40. Farina D, Negro F, Dideriksen JL (2014a) The effective neural drive to muscles is the common synaptic input to motor neurons. The Journal of physiology 592:3427–3441.

41. Farina D, Merletti R, Enoka RM (2004) The extraction of neural strategies from the surface EMG. J.Appl.Physiol. 96:1486–1495.

42. Fuglevand AJ, Winter DA, Patla AE (1993) Models of recruitment and rate coding organization in motor-unit pools. Journal of neurophysiology 70:2470–2488.

43. Gallina A, Disselhorst-Klug C, Farina D, Merletti R, Besomi M, Holobar A, Enoka RM, Hug F, Falla D, Søgaard K (2022) Consensus for experimental design in electromyography (CEDE) project: High-density surface electromyography matrix. Journal of Electromyography and Kinesiology 64:102656.

44. Glaser V, Holobar A. Motor unit identification from high-density surface electromyograms in repeated dynamic muscle contractions. IEEE Transactions on Neural Systems and Rehabilitation Engineering 2018, 27:66–75. pmid:30571641

45. Gogeascoechea A, Ornelas-Kobayashi R, Yavuz US, Sartori M (2023) Characterization of motor unit firing and twitch properties for decoding musculoskeletal force in the human ankle joint in vivo. IEEE Transactions on Neural Systems and Rehabilitation Engineering .

46. Hatze H (1980) Myocybernetic Control Models of Skeletal Muscle - Characteristics and Applications. University of South Africa Press, Pretoria, .

47. Hatze H (1978) A general myocybernetic control model of skeletal muscle. Biological Cybernetics 28 143–157. Heckman, C. J. & Enoka, R. M. (2004) Physiology of the motor neuron and the motor unit. In Anonymous *Handbook of Clinical Neurophysiology.* Elsevier, pp. 119-147.

48. Heckman CJ, Enoka RM (2012) Motor unit. Compr. Physiol. 2:2629–2682.

49. Henneman E (1981) Recruitment of motoneurons : The size principle. Motor Unit Types, Recruitment and Plasticity in Health and Disease 26–60.

50. Henneman E (1957) Relation between size of neurons and their susceptibility to discharge. Science 126:1345–1347.

51. Holobar A, Farina D (2014) Blind source identification from the multichannel surface electromyogram. PM 35:R143-R165.

52. Hug F, Avrillon S, Ibáñez J, Farina D (2023) Common synaptic input, synergies and size principle: Control of spinal motor neurons for movement generation. J.Physiol.(Lond.) 601:11–20.

53. Hug F, Avrillon S, Del Vecchio A, Casolo A, Ibanez J, Nuccio S, Rossato J, Holobar A, Farina D (2021a) Analysis of motor unit spike trains estimated from high-density surface electromyography is highly reliable across operators. Journal of Electromyography and Kinesiology 58:102548.

54. Hug F, Del Vecchio A, Avrillon S, Farina D, Tucker K (2021b) Muscles from the same muscle group do not necessarily share common drive: Evidence from the human triceps surae. J.Appl.Physiol. 130:342–354.

55. Jenz ST, Beauchamp JA, Gomes MM, Negro F, Heckman CJ, Pearcey GE (2023) Estimates of persistent inward currents in lower limb motoneurons are larger in females than in males. J.Neurophysiol. 129:1322–1333.

56. Jung MK, Muceli S, Rodrigues C, Megía-García Á, Pascual-Valdunciel A, Del-Ama AJ, Gil-Agudo A, Moreno JC, Barroso FO, Pons JL (2021) Intramuscular EMG-driven musculoskeletal modelling: Towards implanted muscle interfacing in spinal cord injury patients. IEEE Transactions on Biomedical Engineering 69:63–74.

57. Kapelner T, Sartori M, Negro F, Farina D (2020) Neuro-musculoskeletal mapping for man-machine interfacing. Scientific reports 10:1–10.

58. Kernell D (2006) The Motoneurone and its Muscle Fibres. Oxford University Press UK, .

59. Lloyd, D. G., Jonkers, I., Delp, S. L., & Modenese, L. (2023). The History and Future of Neuromusculoskeletal Biomechanics. Journal of Applied Biomechanics, 39(5), 273–283.

60. Martinez-Valdes E, Enoka RM, Holobar A, McGill K, Farina D, Besomi M, Hug F, Falla D, Carson RG, Clancy EA (2023) Consensus for experimental design in electromyography (CEDE) project: Single motor unit matrix. Journal of Electromyography and Kinesiology 68:102726.

61. Merletti R, Cerone GL (2020) Tutorial. surface EMG detection, conditioning and pre-processing: Best practices. Journal of Electromyography and Kinesiology 54:102440.

62. Merletti R, Muceli S (2019) Tutorial. surface EMG detection in space and time: Best practices. Journal of Electromyography and Kinesiology 49:102363.

63. Muceli S, Poppendieck W, Holobar A, Gandevia S, Liebetanz D, Farina D (2022) Blind identification of the spinal cord output in humans with high-density electrode arrays implanted in muscles. Science advances 8:eabo5040.

64. Muceli S, Poppendieck W, Negro F, Yoshida K, Hoffmann KP, Butler JE, Gandevia SC, Farina D (2015) Accurate and representative decoding of the neural drive to muscles in humans with multi-channel intramuscular thin-film electrodes. J.Physiol.(Lond.) 593:3789–3804.

65. Nawab, Chang, De Luca, High-yield decomposition of surface EMG signals. Clin Neurophysiol 121, 1602–1615 (2010).

66. Negro F, Muceli S, Castronovo AM, Holobar A, Farina D (2016) Multi-channel intramuscular and surface EMG decomposition by convolutive blind source separation. JNE 13:026027.

67. Negro F, Farina D (2011) Decorrelation of cortical inputs and motoneuron output. Journal of Neurophysiology 106:2688–2697.

68. Oliveira AS, Negro F. Neural control of matched motor units during muscle shortening and lengthening at increasing velocities. J.Appl.Physiol. 2021, 130:1798–1813. pmid:33955258

69. Ornelas-Kobayashi R, Gogeascoechea A, Sartori M (2023) Person-specific biophysical modeling of alpha- motoneuron pools driven by in vivo decoded neural synaptic input. IEEE transactions on neural systems and rehabilitation engineering 31:1532–1541.

70. Potvin JR, Fuglevand AJ (2017) A motor unit-based model of muscle fatigue. PLoS computational biology 13:e1005581.

71. Powers RK, Binder MD (2001) Input-output functions of mammalian motoneurons. Rev. Physiol. Biochem. Pharmacol. 143:137–263.

72. Powers RK, Heckman CJ (2017) Synaptic control of the shape of the motoneuron pool input-output function. J.Neurophysiol. 117:1171–1184.

73. Puttaraksa G, Muceli S, Barsakcioglu DY, Holobar A, Clarke AK, Charles SK, Pons JL, Farina D (2022) Online tracking of the phase difference between neural drives to antagonist muscle pairs in essential tremor patients. IEEE Transactions on Neural Systems and Rehabilitation Engineering 30:709–718.

74. Raikova R, Celichowski J, Angelova S, Krutki P (2018) A model of the rat medial gastrocnemius muscle based on inputs to motoneurons and on an algorithm for prediction of the motor unit force. Journal of neurophysiology 120:1973–1987.

75. Rossato J, Hug F, Tucker K, Lacourpaille L, Farina D, Avrillon S (2023) I-spin live: An open-source software based on blind-source separation for decoding the activity of spinal alpha motor neurons in real-time. bioRxiv 2023.04. 14.536933.

76. Sartori M, Yavuz U, Farina D (2017) In vivo neuromechanics: Decoding causal motor neuron behavior with resulting musculoskeletal function. Scientific reports 7:13465–14.

77. Sartori M, Reggiani M, Farina D, Lloyd DG (2012) EMG-driven forward-dynamic estimation of muscle force and joint moment about multiple degrees of freedom in the human lower extremity. PloS one 7:e52618.

78. Teeter C, Iyer R, Menon V, Gouwens N, Feng D, Berg J, Szafer A, Cain N, Zeng H, Hawrylycz M, Koch C, Mihalas S (2018) Generalized leaky integrate-and-fire models classify multiple neuron types. Nat Commun 9:1–15.

79. Thompson CK, Negro F, Johnson MD, Holmes MR, McPherson LM, Powers RK, Farina D, Heckman CJ (2018) Robust and accurate decoding of motoneuron behaviour and prediction of the resulting force output.J.Physiol.(Lond.) 596:2643–2659.

80. Valli G, Ritsche P, Casolo A, Negro F, De Vito G (2024) Tutorial: Analysis of central and peripheral motor unit properties from decomposed high-density surface EMG signals with openhdemg. Journal of Electromyography and Kinesiology 74:102850.

81. Watanabe RN, Magalhães FH, Elias LA, Chaud VM, Mello EM, Kohn AF (2013) Influences of premotoneuronal command statistics on the scaling of motor output variability during isometric plantar flexion. Journal of Neurophysiology 110:2592–2606.

82. Yeung D, Negro F, Vujaklija I (2023) Optimal motor unit subset selection for accurate motor intention decoding: Towards dexterous real-time interfacing. IEEE Transactions on Neural Systems and Rehabilitation Engineering.

